# Biologically plausible unsupervised learning in neural networks with sparse and asymmetric connectivity

**DOI:** 10.1101/2022.11.30.518534

**Authors:** Paul J N Brodersen, Colin J Akerman

## Abstract

In the search for biologically plausible but mathematically precise theories of learning in the brain, recent studies have begun to investigate how key assumptions underlying algorithms for supervised learning in artificial neural networks can be relaxed in biologically plausible ways. Turning to unsupervised learning, we develop biologically more plausible variants of the restricted Boltzmann machine (RBM), and benchmark their performance on MNIST. We show that RBMs with asymmetric connectivity can still be successfully trained with contrastive divergence, even if no two units are reciprocally connected. Furthermore, RBMs are able to learn if the forward, visible-to-hidden layer weights are kept constant and only the backward, hidden-to-visible layer weights are updated. These findings indicate that neural networks with biologically plausible connectivity support contrastive learning.

## 1 Introduction

A restricted Boltzmann machine (RBM) is an artificial neural network that, in its simplest form, consists of two layers of Boltzmann units. A Boltzmann unit is a stochastic binary unit that is active with a probability that is equal to the logistic function of the sum of its connection weights to other active units and a bias term. The connectivity in the network is limited (“restricted”) to connections between units from different layers. The activity of a unit is hence independent of the activity of other units within the same layer, and depends only on the activities of units in the other layer. RBMs were originally developed by Paul Smolensky as a way to learn a generative graphical model that abstracts higher order, latent features from a set of training inputs (Smolensky, 1986). In this model, the input features are represented by the units in the so-called “visible” layer. The higher order, latent features are represented by units in the “hidden” layer. The presence or absence of a feature is represented by the activity of the corresponding Boltzmann unit, which is either “ON” or “OFF”. The central idea of the RBM algorithm is to learn a set of weights for the connections between the two layers such that the hidden unit activity patterns are consistent with visible unit activity patterns for any given input. This consistency is achieved if, for a set of inputs, the visible unit activities result in hidden unit activities that on the next iteration reconstruct the visible unit activities again. To reflect this consistency criterion, the RBM was originally referred to as a “Harmonium”. As a consequence, RBMs not only learn to represent latent features in the data but are also a type of auto-encoder that can be used to reconstruct noisy or partial inputs. The algorithm used in training these networks is called contrastive Hebbian learning (CHL).

Although RBMs were initially envisaged as networks of stochastic binary units, it was quickly discovered that the learning rules could in principle be applied to train networks of deterministic units with linear and logistic activation functions (Peterson and Anderson, 1987), so-called deterministic Boltzmann machines (DBMs). Furthermore, these DBMs could be trained to perform classification and regression tasks, and it was shown that in the regime of weak feedback from the output and upper hidden layers to the lower layers, the weight updates prescribed by contrastive Hebbian learning approximate the updates prescribed by backpropagation of errors (Hinton and McClelland, 1987; O’Reilly, 1996; Xie and Seung, 2003). Subsequently, Geoff Hinton and colleagues discovered a fast variant of the original contrastive Hebbian learning algorithm (contrastive divergence (Hinton, 2002; Carreira-Perpiñán and Hinton, 2005)) such that RBMs could be applied to complex, non-trivial inputs. They showed that the hidden units would learn to encode higher order features (e.g. the strokes in images of handwritten digits in the MNIST data set) and they demonstrated that the activity patterns in small hidden layers could form well-separated, compressed representations of the corresponding input patterns (Hinton and Salakhutdinov, 2006). As a consequence, stacks of pre-trained RBMs form good initialisations for deep artificial neural networks with multiple layers of hidden units that are then trained in classification or regression tasks using backpropation of errors (“backprop”) (Rumelhart et al., 1986). Transporting error gradients in these networks is initially often difficult, which significantly slows gradient descent. Hinton and colleagues showed that pre-training with the RBM algorithm significantly improved performance of these networks (Hinton and Osindero, 2006; Hinton and Salakhutdinov, 2006). Interestingly, the fine tuning of the hidden layer connectivity with backprop turned out to be comparatively small (Erhan et al., 2010).

Although artificial neural networks trained with the RBM algorithm or the backprop algorithm were originally envisioned as models for biological neuronal networks, both algorithms postulate a symmetric connectivity between neurons. In an RBM, the connection from unit *i* to unit *j* is identical to (and has the same weight as) the connection from unit *j* to unit *i*. In back-prop, errors with respect to the activity of a neuron *i* are backpropagated to the neuron *j* in the layer below, proportional to the weight of the (forward) connection from *j* to *i*. This symmetry in the connectivity is in contrast to biological neuronal networks, where the vast majority of synapses transmit information in one direction (with the exception of electrical gap junctions), and only a small percentage of neurons appear bi-directionally connected to each other. Recently, Lillicrap *et al*. have shown that for backprop this symmetry constraint can be abolished (Lillicrap et al., 2016). In their back-prop variant called “feedback alignment”, successive layers are connected by distinct forward and backward weights. The forward weights are learnt by backpropagating errors arising at the output layer via the distinct backward weights to the lower layers. The randomly initialised backward weights are kept fixed for the entirety of training. Despite the random feedback error, such a network can rapidly learn to perform the task as well as a network trained with the canonical backpropation algorithm. Analogously, Detorakis et al. (2019) have shown that a network of deterministic logistic units can be trained using constrastive Hebbian learning with fixed random feedback matrices (rCHL) in the regime of weak feedback from upper layers to lower layers, i.ew? hen contrastive Hebbian learning approximates backpropagation of errors.

However, in biological neuronal networks, activity in sensory cortical areas can be driven by top-down activity. As biological neurons have a spike threshold, this top-down input necessarily is of similar magnitude as the bottom up activity. Clearly, in biological neuronal networks recurrent feedback connections can be strong. Here, we hence investigate whether the symmetry constraint on the connectivity can similarly be relaxed in an RBM consisting of stochastic binary units, where feedback connections are on average at least as strong as the forward connections. We show that this is the case in two different ways: First, we demonstrate that an RBM with distinctly initialised forward and backward weights can still successfully learn the task. However, as the same weight updates are applied to the forward and backward weights, one might argue that this is unsurprising as forward and backward weights can become sufficiently similar with learning as long as the cumulative weight changes are sufficiently large. We hence demonstrate that this finding remains true in a sparse network that does not have any bi-directional connections. For Boltzmann machines with deterministic units, this has previously been noted in Galland and Hinton (1991). Second, we show that updates only to the backward weights suffice to learn to reconstruct a set of inputs. This is reminiscent of feedback alignment and rCHL, only that in this case the forward weights are kept constant and the backward weights are learnt. These findings have significant implications for the development of novel RBM variants and for the plausibility of RBM-like networks as models for biological neuronal networks.

## 2 Methods

### 2.1 Data acquisition and pre-processing

All experiments were carried out using images from the MNIST database of handwritten digits, which was retrieved from http://yann.lecun.com/exdb/mnist/. The original train and test split of the data set was retained. Example inputs and example reconstructions correspond to the first 100 samples in the test data set. As the states of the visible units are constrained to be between 0 and 1, all inputs were normalised to this range by dividing image pixel intensities by 255.

### 2.2 The restricted Boltzmann machine

All RBM networks consisted of two layers: a visible layer with 784 units corresponding to the 784 pixels in a single MNIST image, and a hidden layer of 400 units, i.e. a layer with about half as many units as in the visible layer. A hidden layer with fewer units (and hence features) than present in the visible layer ensures that the transformation learnt by the RBM does not simply correspond to the identity transform, and demonstrates the ability of the RBM to compress the set of input features to a set of hidden features.

All hidden layer units were modelled as stochastic Boltzmann units. A stochastic Boltzmann unit is in the active state with the probability given by the logistic function of the sum of its inputs from visible units and a bias term:

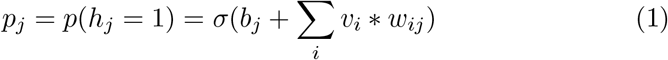

Here, *i* and *j* index the visible units *v*_*i*_ and hidden units *h*_*j*_, respectively, *b* is the bias term associated with each unit, and *w*_*i*_*j* is the weight of the connection from *v*_*i*_ to *h*_*j*_. The function ∼ represents the logistic function, which is defined as:

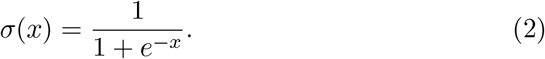

Visible layer units were modelled similarly:

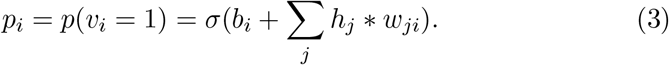

However, unlike hidden layer units that transmitted their binary state to visible units, visible units transmitted the probability of being active *p*_*i*_ instead of their state *v*_*i*_. Such a procedure has previously been suggested to reduce sampling noise and thus speed up learning (Hinton, 2012). Strictly speaking, the visible units were hence modelled as logistic units, and not as Boltzmann units.

### 2.3 Weight and bias updates

Weights and biases were updated according to the canonical RBM update rules:

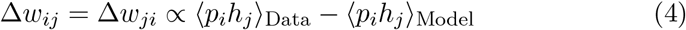

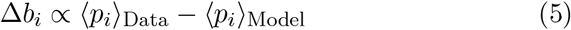

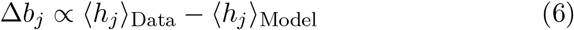

As before, *p*_*i*_ is the probability that the visible unit *i* is active, *h*_*j*_ is the activity state of hidden unit *j*, and *w*_*ij*_ is the weight of the connection from *i* to *j*. The term ⟨*p*_*i*_*h*_*j*_⟩ with the subscript “Data” thus corresponds to the co-activity state of visible unit *i* and hidden unit *j* when the visible units are clamped at the input sample values. The subscript “Model” indicates the co-activity state when the visible units are not clamped to a predetermined value. The “Data” and “Model” activity states were sampled using contrastive divergence. Specifically, the RBM was initialised with the visible unit states set to the values of an input sample, and on that basis the hidden unit states were computed. These unit states formed the basis of the “Data” samples. To sample the “Model” states, the network was allowed to evolve for a number of *CD* additional backward and forward passes of the activity:

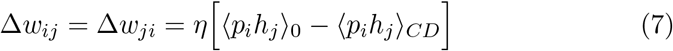

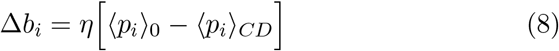

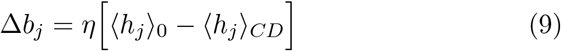

All experiments were performed with *CD* = 3 contrastive divergence iterations, and a learning rate *η* = 0.01.

### 2.4 Weight and bias initialisation

Unless stated otherwise, biases were initialised with a value of 0. All weights were initialised with values drawn from the normal distribution *N* (0.0, 0.1).

### 2.5 Software implementation

The code to reproduce all experiments shown in the figures below can be accessed at https://github.com/paulbrodersen/rbm_variants. The MNIST data was loaded into memory using the python-mnist package. All RBM variants were implemented in python. Numerical computations relied in part on functions from the numpy package. All visualisation were made using the matplotlib package.

## 3 Results

### 3.1 Restricted Boltzmann machines with independent and asymmetric forward and backward weights learn to reconstruct inputs

A canonical RBM forms an undirected graph: if neuron *i* is connected to neuron *j* with weight *w*_*ij*_, then neuron *j* is connected to neuron *i* with weight *w*_*ji*_ = *w*_*ij*_. We wanted to determine whether network architectures that are less constrained in their connectivity are still able to learn and perform the same computation. As a first pass, we evaluated the performance of RBMs with independently initialised forward and backward weights (i.e. *w*_*ij*_ ≠ *w*_*ji*_) on a standard machine learning data set, the MNIST database of handwritten digits. In analogy to directed graphs, we call such an RBM a “directed” RBM. Fig. 3.1 compares the performance of a canonical, undirected RBM network with the performance of such a directed RBM with distinct forward and backward weights. The directed RBM is still clearly able to learn to reconstruct the MNIST digits, albeit with a somewhat lower performance.enHidden

**Figure 1:**
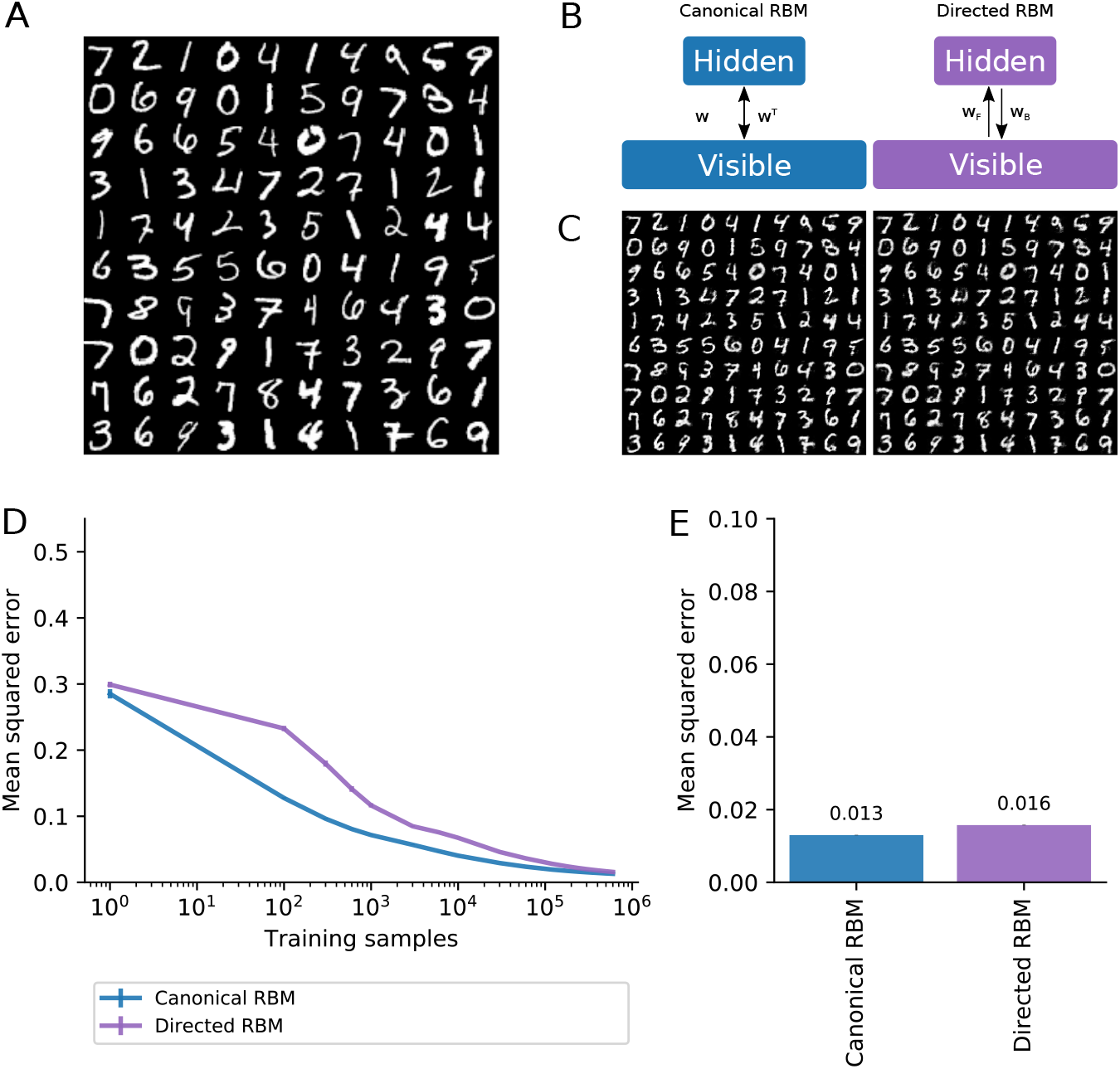
Distinct, asymmetric forward and backward weights are sufficient for learning. A canonical RBM and a directed RBM were trained to reconstruct images in the MNIST handwritten digits database. Each RBM consisted of 784 visible units and 400 hidden units. Biases and connection weights were updated using contrastive divergence with 3 iterations and a learning rate *η* = 0.01. a) Images of the first 100 characters in the MNIST test set. b) Cartoon representation of a canonical RBM (left), in which the backward weight matrix *w*^*T*^ is the transpose of the forward weight matrix *w*. Cartoon representation of a directed RBM (right), in which the forward weight matrix *w*_*F*_ is distinct of the backward weight matrix *w*_*B*_. c) Example reconstructions of the first 100 characters in the test set after 10 epochs of learning on the training set by a canonical RBM (left) and a directed RBm (right). d) Mean squared error between the input images and RBM reconstructions for the test set plotted against the number of training samples. e) Final mean squared error after 10 epochs. Error bars represent the standard deviation around the mean (*n* = 10).

**Figure 2:**
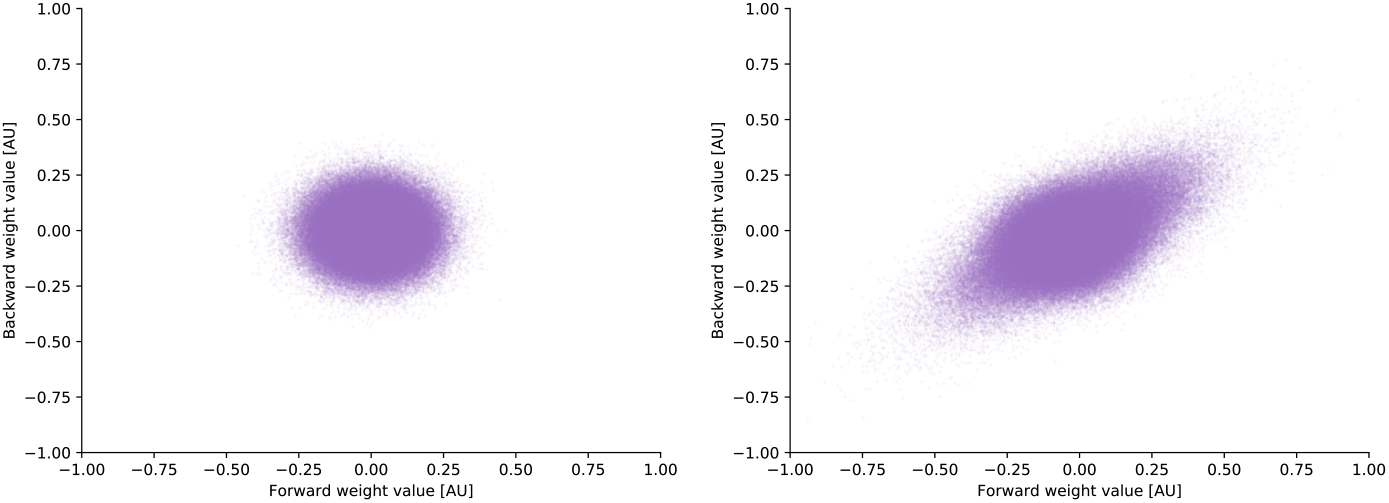
Forward and backward weights in directed RBMs become more similar with training. Forward weights *w*_*ij*_ plotted against the corresponding backward weights *W*_*ji*_ before and after training a directed RBM network for 10 epochs on MNIST.

**Figure 3:**
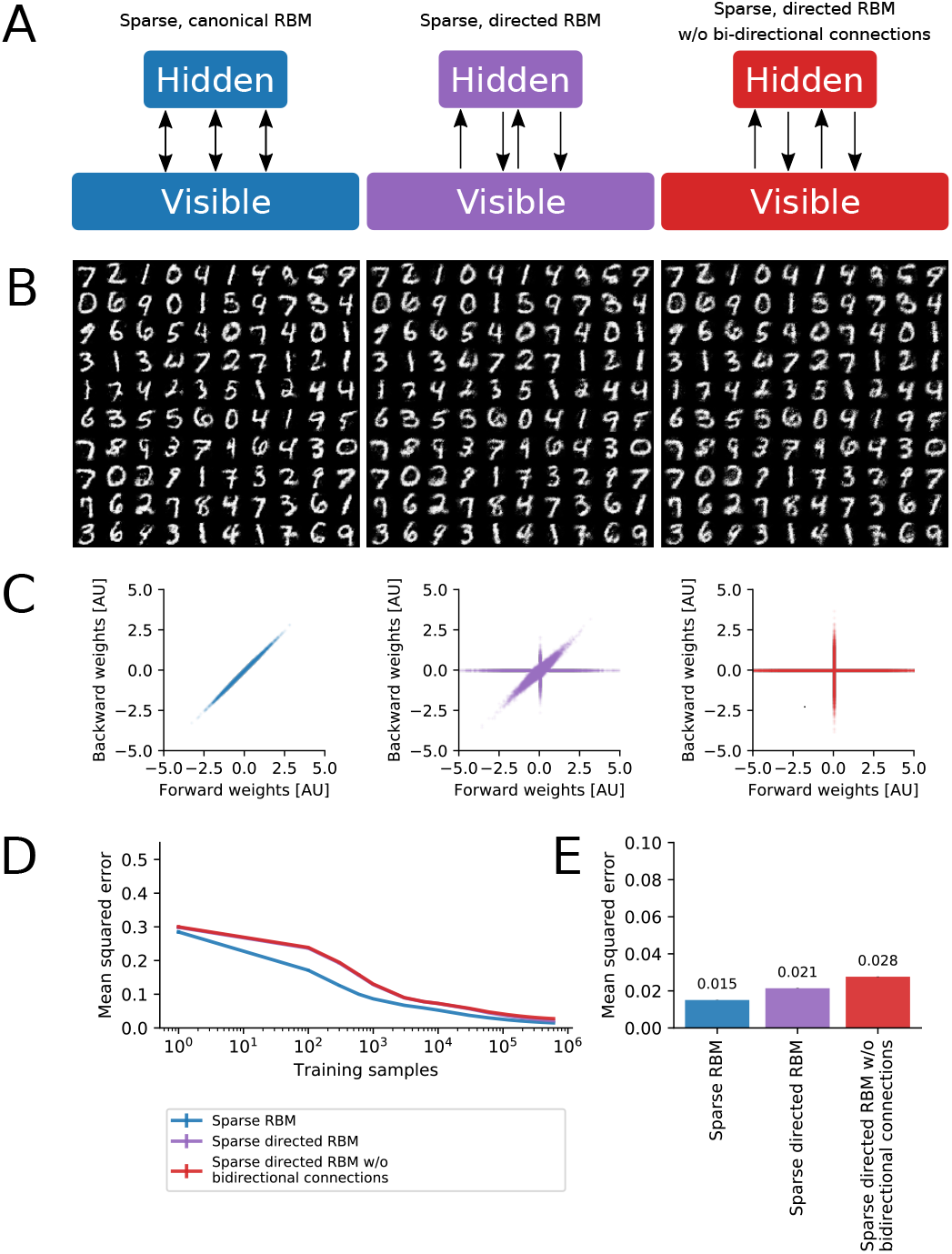
Uni-directional connections are sufficient for learning: Three sparsely connected RBM variants with differing proportions of bi-directional connections were trained for 10 epochs on the MNIST handwritten digit data set. As before, each RBM consisted of 784 visible units and 400 hidden units, and biases and connection weights were updated using contrastive divergence with 3 iterations and a learning rate *η* = 0.01. However, the connectivity between units in different layers was sparse (*p* = 0.5), and the three variants differed in the proportion of bi-directional connections (*w*_*ij*_ ≠ 0*∧* w_*ji*_ ≠ 0), which was set to 1.0 (first column), 0.25 (second column), and 0.0 (third column). a) Cartoon representation of the sparse RBM variants tested. b) Corresponding example reconstructions of the first 100 characters in the test set after 10 epochs of learning on the training set. c) Corresponding size of the forward weights *w*_*ij*_ versus the corresponding backward weights *w*_*ji*_ for an example network after training for 10 epochs. d) Mean squared error between the input images and RBM reconstructions for the test set plotted against the number of training samples. e) Final mean squared error after 10 epochs. Error bars correspond to the standard deviation around the mean (*n* = 10).

However, as we did not change the weight update rules for the directed RBMs, the same weight updates were applied to the forward and backward weights. As sufficiently large weight changes accumulate over time, the forward and backward weight matrices become more similar to one another. This effect can be seen in Fig. 3.1, which plots forward weights against the respective backward weights before and after training. The forward and backward weights clearly align to one another with learning, as indicated by the elongation of the data along the line of equality, i.e. where *y* = *x*.

Although the weight alignment tends to be small for sensibly initialised network architectures, the alignment could drive learning in the directed RBMs by effectively reducing the directed networks to approximate undirected networks. To show that this was not the case, we compared three sparsely connected RBM variants, namely (1) a canonical RBM (i.e. with purely bi-directional connections), (2) a directed RBM with independently initialised forward and backward connections containing a mix of bi-directional and uni-directional connections, and (3) a directed RBM constrained to have purely uni-directional connections, i.e. lacking any reciprocal connections. As can be seen in Fig. 3.1, RBM performance decreased with a reduction in the number of bi-directional connections. However, the absolute performance decrease from an RBM with purely bi-directional connections, to an RBM with purely uni-directional connections, was overall very small, at 1.3%. These findings show that weight alignment was not necessary for learning, but that learning uni-directional connections was sufficient for training an RBM. Perfect symmetry in the forward and backward connectivity thus may confer some advantage in learning, but is not necessary.

### 3.2 Updating the backward weights is sufficient for learning

In the previous section, we showed that learning uni-directional connections is sufficient to train an RBM. Lillicrap *et al*. previously showed that neural networks can be trained with backpropagation of errors via fixed backward weights (Lillicrap et al., 2016). By analogy, we hypothesised that RBMs could also be trained by updating only one set of connections, i.e. either the forward or the backward weights. To test this hypothesis, we compared five RBM variants: (1) the canonical RBM network, (2) a directed RBM, where both the forward and the backward weights were updated, 3) a directed RBM where the forward weights were kept fixed and the backward weights were updated as before, 4) a directed RBM where the forward weights were updated and the backward weights were kept fixed, and (5) a directed RBM where both forward and backward weights were kept fixed (and only the unit biases were updated). As shown in Fig. 4, updating only the backward weights was sufficient for learning to reconstruct the inputs with a performance that was comparable to (albeit slightly smaller than) the performance of a canonical RBM. Interestingly, the “inverse” variant, i.e. learning the forward weights while keeping the backward weights fixed, also performed better than just learning the biases, although the performance of this RBM variant was substantially lower than in the canonical case.id

**Figure 4:**
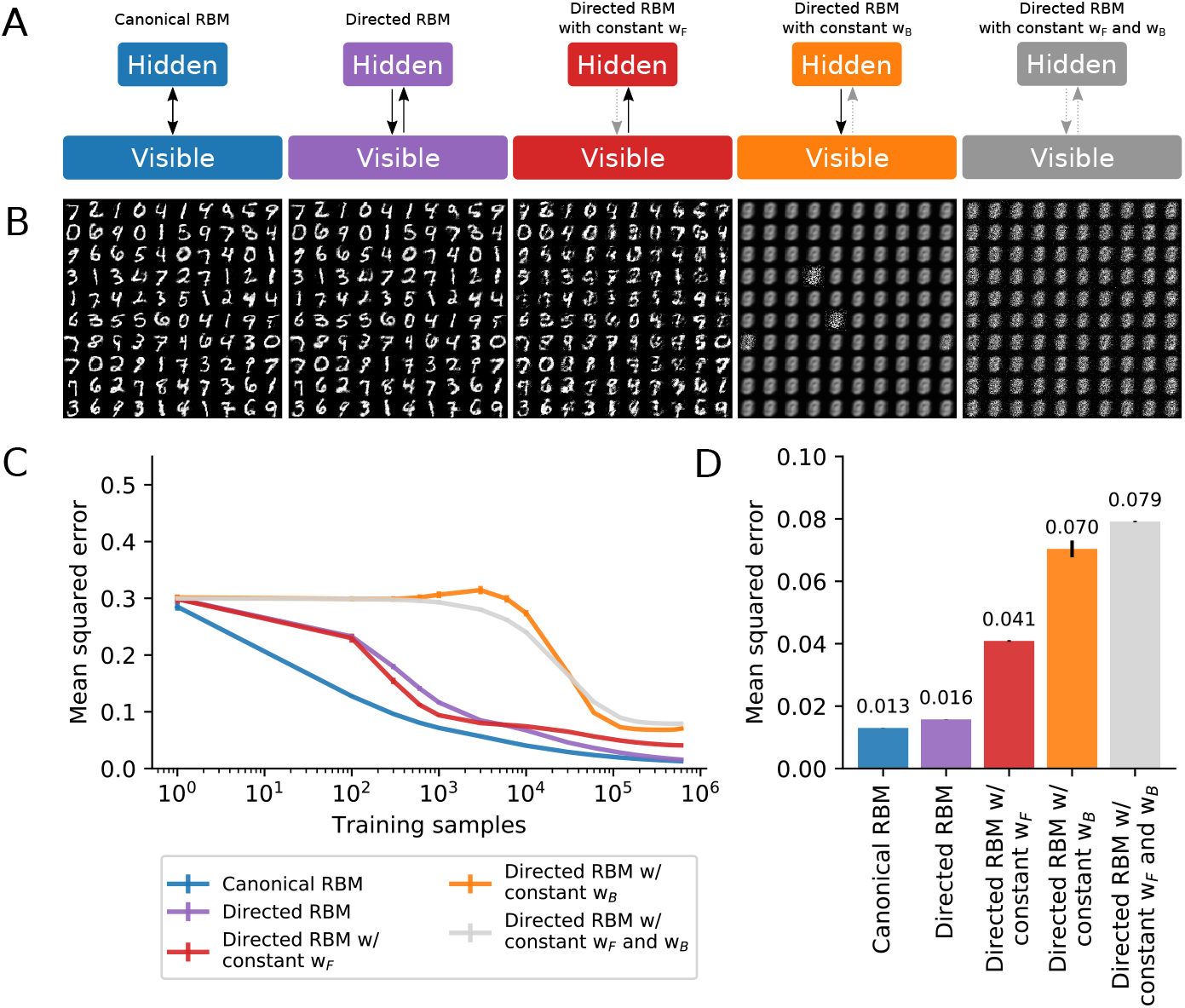
Updating the backward weights is sufficient for learning: A canonical RBM and four directed RBM variants were trained for 10 epochs on the MNIST hand-written digit data set. As before, each RBM consisted of 784 visible units and 400 hidden units, and biases and connection weights were updated using contrastive divergence with 3 iterations and a learning rate *η* = 0.01.a) Cartoon representation of RBM variants tested (in order): a canonical RBM, a directed RBM, where forward weights, backward weights, and unit biases were updated, a directed RBM, where the forward weights were kept fixed, a directed RBM, where the backward weights were kept fixed, and a directed RBM, where both sets of weights were kept fixed and only the unit biases were updated. b) The corresponding example reconstructions of the first 100 characters in the test set after 10 epochs of learning on the training set. c) Mean squared error between the input images and RBM reconstructions for the test set plotted against the number of training samples. d) Final mean squared error after 10 epochs. Error bars correspond to the standard deviation around the mean (*n* = 10).

## 4 Discussion

Here we establish that the symmetry constraint imposed on the forward and backward weights in the canonical RBM is not necessary for learning. Instead, even networks with no reciprocal connections are still able to learn, as are networks in which only the backward weights are updated. These findings indicate that there is a family of network architectures with diverse connectivity schemes that support RBM-like learning. This diversity, in turn, opens up the possibility that some biological neuronal networks may learn to extract features from their sensory inputs using contrastive learning. Nevertheless, it is worth noting that the canonical RBM outperforms all other variants explored here. This indicates that bi-directional, symmetric connections may bequeath a fundamental advantage for learning certain relationships. Interestingly, bi-directional connections, whilst rare, seem to be more common in the brain than expected by chance (Song et al., 2005; Perin et al., 2011), and the findings reported here might point towards reasons why that is the case.

## References

Carreira-Perpiñán, M. a. and Hinton, G. E. (2005). On Contrastive Divergence Learning. Artificial Intelligence and Statisticss, 0:17.

Detorakis, G., Bartley, T., and Neftci, E. (2019). Contrastive Hebbian learning with random feedback weights. Neural Networks, 114:1–14.

Erhan, D., Courville, A., and Vincent, P. (2010). Why Does Unsupervised Pre-training Help Deep Learning? Journal of Machine Learning Research, 11:625–660.

Galland, C. C. and Hinton, G. E. (1991). Deterministic Boltzmann Learning in Networks with Asymmetric Connectivity. In Connectionist Models, pages 3–9. Elsevier.

Hinton, G. and Osindero, S. (2006). A fast learning algorithm for deep belief nets. Neural computation, 1554:1527–1554.

Hinton, G. E. (2002). Training products of experts by minimizing contrastive divergence. Neural Computation, 14(8):1771–1800.

Hinton, G. E. (2012). A Practical Guide to Training Restricted Boltzmann Machines, pages 599–619. Springer Berlin Heidelberg, Berlin, Heidelberg.

Hinton, G. E. and McClelland, J. (1987). Learning representations by recirculation. In Anderson, D., editor, Neural Information Processing Systems, volume 0. American Institute of Physics.

Hinton, G. E. and Salakhutdinov, R. R. (2006). Reducing the Dimensionality of Data with Neural Networks. Science, 313(5786):504–507.

Lillicrap, T. P., Cownden, D., Tweed, D. B., and Akerman, C. J. (2016). Random synaptic feedback weights support error backpropagation for deep learning. Nature Communications, 7:1–10.

O’Reilly, R. C. (1996). Biologically Plausible Error-Driven Learning Using Local Activation Differences: The Generalized Recirculation Algorithm. Neural Computation, 8(5):895–938.

Perin, R., Berger, T. K., and Markram, H. (2011). A synaptic organizing principle for cortical neuronal groups. Proceedings of the National Academy of Sciences of the United States of America, 108(13):5419–5424.

Peterson, C. and Anderson, J. (1987). A mean field theory learning algorithm for neural networks. Complex Systems, 1:995–1019.

Rumelhart, D. E., Hinton, G. E., and Williams, R. (1986). Learning representations by back-propagating errors. Nature, 323(9).

Smolensky, P. (1986). Information processing in dynamical systems: Foundations of harmony theory. Parallel Distributed Processing Explorations in the Microstructure of Cognition, 1(1):194–281.

Song, S., Sjöström, P. J., Reigl, M., Nelson, S., and Chklovskii, D. B. (2005). Highly nonrandom features of synaptic connectivity in local cortical circuits. PLoS biology, 3(3):e68.

Xie, X. and Seung, H. S. (2003). Equivalence of Backpropagation and Contrastive Hebbian Learning in a Layered Network. Neural Computation, 15(2):441–454.

